# Machine learning to predict microbial community functions: An analysis of dissolved organic carbon from litter decomposition

**DOI:** 10.1101/599704

**Authors:** Jaron Thompson, Renee Johansen, John Dunbar, Brian Munsky

**Author notes:** Corresponding author: (BM).

## Abstract

Microbial communities are ubiquitous and often influence macroscopic properties of the ecosystems they inhabit. However, deciphering the functional relationship between specific microbes and ecosystem properties is an ongoing challenge owing to the complexity of the communities. This challenge can be addressed, in part, by integrating the advances in DNA sequencing technology with computational approaches like machine learning. Although machine learning techniques have been applied to microbiome data, use of these techniques remains rare, and user-friendly platforms to implement such techniques are not widely available. We developed a tool that implements neural network and random forest models to perform regression and feature selection tasks on microbiome data. In this study, we applied the tool to analyze soil microbiome (16S rRNA gene profiles) and dissolved organic carbon (DOC) data from a 44-day plant litter decomposition experiment. The microbiome data includes 1709 total bacterial operational taxonomic units (OTU) from 300+ microcosms. Regression analysis of predicted and actual DOC for a held-out test set of 51 samples yield Pearson’s correlation coefficients of .636 and .676 for neural network and random forest approaches, respectively. Important taxa identified by the machine learning techniques are compared to results from a standard tool (indicator species analysis) widely used by microbial ecologists. Of 1709 bacterial taxa, indicator species analysis identified 285 taxa as significant determinants of DOC concentration. Of the top 285 ranked features determined by machine learning methods, a subset of 86 taxa are common to all feature selection techniques. Using this subset of features, prediction results for random permutations of the data set are at least equally accurate compared to predictions determined using the entire feature set. Our results suggest that integration of multiple methods can aid identification of a robust subset of taxa within complex communities that may drive specific functional outcomes of interest.

## Introduction

Microbial communities mediate essential functions in diverse ecosystems. While the microbiome controls many interesting macroscopic properties, elucidating the relationship between specific microbes and ecosystem functions remains a complex problem in ecology. Recent advances in DNA sequencing technology make it easy to acquire metagenomic data representing the taxonomic profile of bacteria and fungi in microbial communities. This opens the door to deciphering which components of the microbiome can drive changes in macroscopic properties. However, analysis of metagenomic microbial data poses several difficulties. The data are typically high dimensional (many taxa) with a small number of samples collected in each study. Additionally, sequencing results are noisy and yield sparse data sets [1].

Machine learning techniques provide a means to analyze high-dimensional data [2,3] and could be used to elucidate relationships between microbial taxa (or other metagenomic features such as gene families or metabolic pathways) and environmental attributes. The random forest model is reportedly one of the most effective machine learning models for analyzing microbiome data; high classification accuracy has been demonstrated with a variety of 16S rRNA data sets for identification of body habitat, host, and disease states [4]. In another study, artificial neural networks were used to map complex relationships between microbial communities and environmental variables, enabling predictions of the abundance of microbial taxa across the English Channel, for example [5].

While most existing machine learning software packages focus on binary classification of microbial data sets [6–8], random forest and neural network models can also be used to identify the subset of microbial taxa whose relative abundances best predict a continuous target variable [9, 10]. The combination of random forest and neural network models can evaluate feature importance and reveal which microbial taxa are most positively or negatively correlated with target variables. To provide helpful perspective for microbial ecologists, we compare results from these machine learning techniques to indicator species analysis, a commonly used tool in ecology that is typically used for classification, though similar techniques have been adapted for regression problems [11]. We also show how our tool can be applied to study the effect of experimental sample size on model performance by evaluating prediction error over increasing subsets of training data. In this study, we apply the proposed random forest and neural network regression models to predict the abundance of dissolved organic carbon (DOC) from plant litter decomposition, where bacterial taxa abundances are treated as model features/variables. Recent studies have shown that microbial communities play an important role in carbon cycling and can potentially be manipulated to increase the abundance of DOC for transport and sequestration in deeper soil layers [12–16]. We use DOC and bacterial community data from a study that examined the role of soil microbial community composition in controlling carbon flow from plant litter decomposition [17]. Feature selection results determined by machine learning methods are compared to indicator analyses [18,19] in which high and low DOC are used as classification category labels. The ultimate goal of this study is to present a powerful set of tools for prediction and feature selection tasks designed specifically for elucidating the relationship between microbial communities and the ecosystems they inhabit.

## Materials and methods

Random forest and neural network regression models are examples of supervised machine learning algorithms. In contrast to unsupervised machine learning algorithms, these methods require a subset of the data called a *training* set to develop a mathematical relationship between *features* and *target* variables. A *feature* represents a model variable and the *target* is the variable the model predicts. For regression problems, the target variable is a continuous scalar, and for classification problems, the target is a discrete label. A *sample* is a single set of features paired with a target variable, which, in the context of the present case study, represents a bacterial community profile paired with DOC. To assess model performance, predicted target variables using features from a held-out set of *test* data are compared to known target variables. In this study, prediction performance is measured using Pearson’s correlation coefficient, which quantifies the linear correlation between predicted and true target variables, and for which a value of one indicates a perfect positive linear correlation. In general, our regression model assumes that targets and features are related to one another by

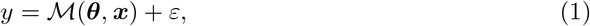

where 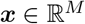 is a vector *M* features, 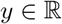 is the corresponding true value of the target variable, 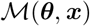 is some mathematical operation (or model) from 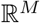 to 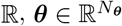 are model parameters, and *ε* is the prediction error.

We denote the set of *M* features with *N* samples as the *N* × *M* feature matrix 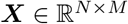, which can be mapped to a vector of *N* target variables 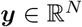 according to

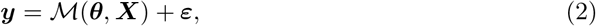

where 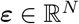 is the vector of prediction errors. While Eq. 2 describes the general regression problem common to most machine learning algorithms, the actual form of 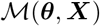 varies according to the specific approach. We introduce a few of these machine learning approaches as follows.

### Neural network regression model

A feed-forward neural network regression model applies a series of parameterized activation functions organized in layers to map features in a sample to a continuous target variable. Each layer of a feed-forward neural network is composed of a set of nodes which apply a nonlinear transformation to the sum of the product of inputs from the previous layer and weight parameters plus an additional bias parameter. A stochastic gradient descent algorithm minimizes the cost function by adjusting model parameters (weights and bias values for each layer) via a process called error back-propagation, which updates model parameters in each layer based on the gradient of the cost function with respect to model parameters. The rate at which model parameters change during training can be adjusted by a learning rate hyper-parameter, and the cost function can be adjusted with a regularization hyper-parameter, which ensures that model parameters do not reach disproportionate values [2]. We built a feed-forward neural network regression model using Theano [20] and Python 3.7 with a randomized search algorithm for determining model hyper-parameters implemented with Scikit-learn [21]. As a default, the model includes a single hidden layer with 15 nodes with sigmoid activation functions and a single output layer with a linear activation function. A randomized hyper-parameter search uses the training data set to find the optimum hidden layer size, learning rate, and regularization coefficient. Our model applies the mean squared error between predicted and true values as a cost function for use with the training and validation analyses. Training the neural network model is an iterative process, where each iteration is called a training epoch. In each training epoch, the total set of training data is divided and trained over randomly chosen mini-batches. Once the cost function applied to the validation data set fails to decrease over a default of ten training epochs, training stops. For this study, the model was trained with 257 training samples and tested with a held-out set of test data with 51 samples. To assess the correlation between true DOC and predicted DOC for each sample, Pearson’s correlation coefficient was computed for training and testing results.

### Neural network feature selection

Methods for evaluating feature importance using a neural network model often focus on weights assigned to individual features after training of the model [22,23]. Our proposed feature selection tool employs a similar approach, where the gradient of the model output with respect to weights associated with each feature is used to determine the feature importance vector. Each element of the feature importance vector corresponds to an individual feature, where the magnitude of each element is indicative of feature importance for predicting the target variable, and the sign indicates whether the feature has a positive or negative impact on the predicted variable.

For a feed-forward neural network model with *M* features as inputs that connect to *J* nodes in the first hidden layer, we can denote the *M × J* matrix of weights connecting each feature to each node as 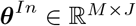, where *θ^In^* is a subset of the full parameter set *θ*. The gradient of the model output with respect to *θ*^In^ provides the *M × J* feature importance matrix, 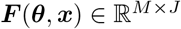, which we define as

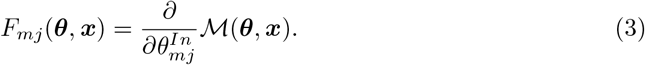

Marginalizing the feature importance matrix over all nodes in the first hidden layer produces a *M*-dimensional vector, which we will call the feature importance vector *f*(*θ*, *x*), whose elements are

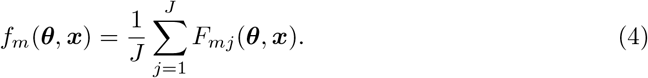

After training the model, we determine the sensitivity of the model to each feature, denoted as the *M*-dimensional vector 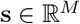, by calculating the average value of the feature importance vector over the set of training data with *K* samples

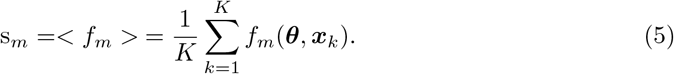

To gain confidence in the importance assigned to features, feature importance is determined using a bootstrap method, which randomly samples 80% of the training data set over a default of 50 iterations. The average feature ranking values determined over all iterations represents the most confident set of ranked features.

### Random forest regression model

Decision tree based machine learning methods map features to target variables by splitting the set of possible target variables based on the values of individual features [2,24]. An *internal node* is a point at which the value of a feature determines a split in the set of possible target variables, and the nodes that follow an internal node are called *leaf nodes* [2]. The random forest method constructs a set of decision trees constructed from randomly selected subsets of the feature space and computes the model output by averaging the predictions from individual decision trees [25]. Using the random forest regressor from Scikit-learn [21], a random forest regression model is instantiated with a mean squared cost function, two samples required to split an internal node, and one sample required to be at a leaf node as the default. Hyper-parameters for the model include the number of samples required to split an internal node, the number of samples required to be at a leaf node, and the number of features to consider in each decision tree. These hyper-parameters can be optimized with the training set using Scikit-learn’s randomized search algorithm. During training, the random forest regressor model fits an ensemble of 1000 decision trees trained on randomly selected sub-samples of the data set. All random forest results from this study use identical training and testing data to allow direct comparison to the neural network model.

### Random forest feature selection

The random forest regressor made available by Scikit-learn [21] returns an array of feature importance values of length equal to the array of input features. Decision tree algorithms, such as random forest, assess feature importance by examining how well a feature (often referred to as variable in literature [24]) can split the potential output labels. In other words, a highly significant feature provides the greatest reduction of potential labels for a given sample. Additionally, feature importance is determined as part of the boot-strap method used for assembling random decision trees, where feature importance is greater for variables that result in greater prediction performance when included in the decision trees [24]. To gain confidence in the rank assigned to features, feature ranking is determined using a bootstrap method that randomly samples 80% of the training data set over a default of 50 iterations. The highest average feature ranking values determined over all iterations represent the most confident ranked features.

### Indicator species analysis for feature selection

Indicator species analysis [18,19] is used for comparison with the feature selection results determined by the above machine learning methods. Indicator species (hereafter we use ‘taxa’, not ‘species’, for accuracy) are defined as the features that are most indicative of changes in DOC across different samples. To determine indicator taxa, a correlation value is calculated for each feature as the product of *specificity* and *fidelity* for a particular taxon in association with either high or low DOC samples [18]. Specificity measures how much a taxon associates with a single label (e.g., high or low DOC), and fidelity measures how frequently a taxon associates with that label. Specificity would be maximized if a taxon were only present in sites with a particular label, and fidelity would be maximized if a taxon were present at all sites associated with a particular label. A confidence score is assigned to each feature using a boot-strap algorithm that compares the correlation value for each feature determined using correct labels with correlations determined using randomly assigned labels. If the correlation statistic between features and site labels determined using random labels is not consistently lower than the correlation statistic using correct labels, then the confidence score for that feature-site correlation is low. Only taxa with at least a 95% confidence (features with correlation values greater than 95% of correlations determined with random labels) are considered in this study. Indicator taxa analysis was implemented in Python 3.7 with the methods described in Dufrene and Legendre, 1997 [18].

### Data acquisition and data pre-processing

Microbiome data (16S rRNA gene profiles) were obtained from a prior study of pine needle litter decomposition in laboratory microcosms [17] (supporting information S1 Dataset). In brief, the microbial community in each of 206 soil samples was suspended in water, inoculated into three replicate microcosms containing sterile sand and pine litter, and incubated 44 days at 25C. At 44 days, the amount of DOC in the microcosms was measured, DNA was extracted from a subset of microcosms, and 16S rRNA gene amplicons were sequenced on an Illumina MiSeq. Details of sequence processing are described in [17]. Sequence data has been deposited in the NCBI sequence read archive (SRP151768). Because the composition of bacterial communities among replicate microcosms diverged over the 44-day incubation period, the replicates were treated as independent samples. For machine learning analysis, however, the training and testing data were prohibited from sharing replicate samples to ensure independence between training and testing data sets (supporting information S2 Dataset, S3 Dataset). The bacterial community profiles from 308 samples were rarefied to 1023 sequences, which yielded a matrix with a total of 1709 bacterial taxa. By default, our tool standardizes features such that each feature is zero mean with unit variance over the training data set. The test data is similarly scaled but only using the sample statistics determined from the training data set.

## Results

Our feed forward neural network regression model was trained with 257 community samples to predict level of DOC (Fig 1A). Our model was tested with a held out set of 51 test samples which yielded a Pearson’s correlation coefficient of .636 between true and predicted DOC (Fig 1B) and a mean squared error of .565. The random forest regression model was trained and tested with identical sets of data used with the neural network model. Test results using the random forest regression model yielded a Pearson’s correlation coefficient of .676 (Fig 1D) and a mean squared error of .516. A scatter plot of the prediction error using the neural network model versus the prediction error with identical test samples using the the random forest model are positively correlated with a Pearson’s correlation coefficient of 0.781 (Fig 1E).

**Fig 1.**
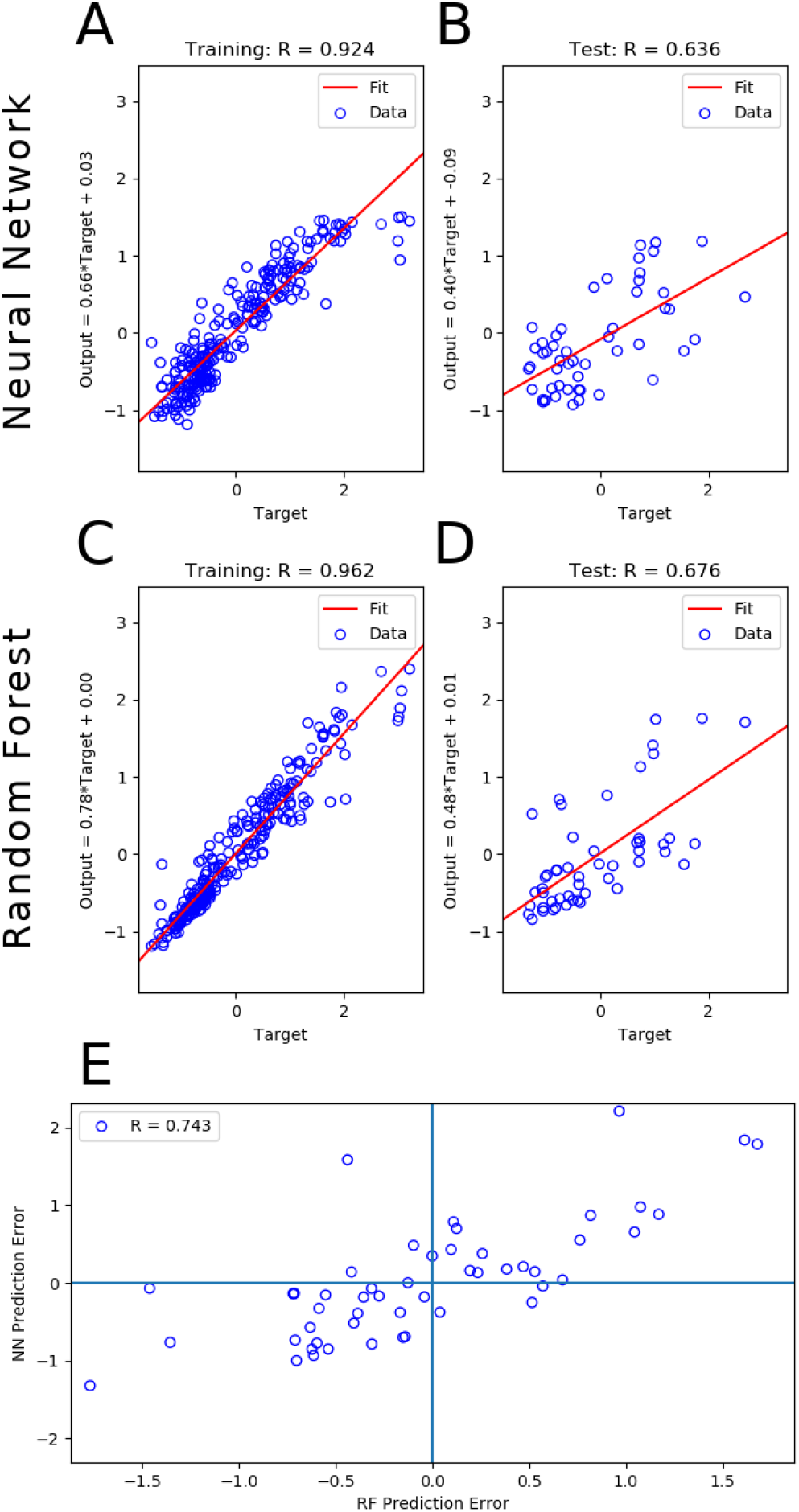
DOC prediction with neural network and random forest regression models. (A) Scatter plot of fitted DOC versus true DOC from training data samples (n=257) using neural network model. (B) Scatter plot of predicted DOC versus true DOC from test data samples (n=51) using neural network model. (C-D) Same as above but using random forest model. Training and testing data are identical for both methods. (E) A scatter plot of the prediction errors using the neural network model versus the prediction errors with identical test samples using the random forest model.

To illustrate the degree of agreement of feature importance for predicting DOC between random forest, neural network, and indicator species approaches, Fig 2A shows a Venn diagram comparing feature selections. Feature selection was performed on the same training set used to produce Fig 1. Out of a feature set with 1709 taxa, 285 taxa were significant indicator taxa. Of the top 285 ranked features from the machine learning methods, 112 bacterial taxa were shared between random forest and neural network feature selections, and of these, 86 bacterial taxa overlapped with the set of indicator taxa. To further investigate agreement of feature importance between methods, Fig 2B shows how the shared set of ranked features determined by the neural network, random forest, and indicator taxa analysis varies as a function of feature rank. To investigate the significance of our feature selection results, we compared the number of features in the consensus set to the number of shared features that would occur if features were selected from three randomly organized sets. We applied a Monte Carlo approach that sampled features from three randomly organized sets of 1709 features and counted the number of features that were commonly selected in a pair of sets or within the intersection of all three sets. We plotted the mean and 99% confidence interval from 1,000 simulations as a function of the number of sampled features (a separate plot with just the Monte Carlo simulation curve is included in the supporting information S4 Fig). The number of features in the consensus set is consistently greater than the number of shared features expected from random sampling, suggesting that each feature selection approach exploited similar, non-random trends in the data. Figs 2A,B show that feature importance determined by the neural network has greater agreement with indicator taxa compared to feature importance determined by random forest.

**Fig 2.**
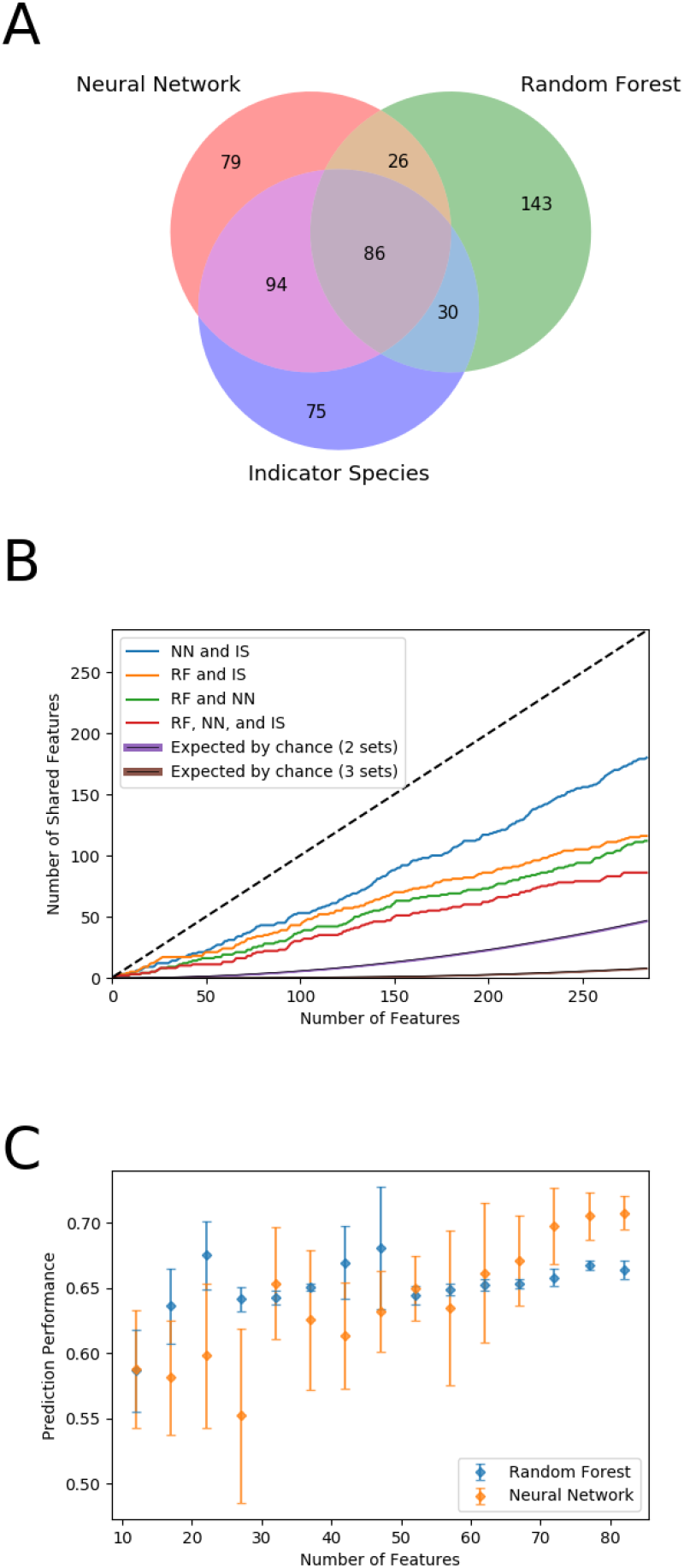
Feature ranking determined by neural network, random forest, and indicator species analysis. (A) Venn diagram demonstrates agreement of 86 bacterial taxa out of the top 285 ranked taxa from machine learning methods. (B) Plots of the number of shared features between NN and IS (blue), RF and IS (orange), RF and NN (green), and all methods (red) as a function feature rank over 285 features. Monte Carlo simulation of the number of shared features expected by randomly sampling from 3 sets of 1709 features is plotted with a 99% confidence interval (black line, purple confidence inteval). The black dotted line indicates perfect agreement between the three sets of ranked features. (C) Plot of prediction performance on test data as measured by Pearson’s correlation coefficient versus number of features included in machine learning models. The data are binned such that each point represents the average prediction over 5 trials, where each subsequent trial includes an additional feature.

Indicator species analysis not only provides a feature importance metric, but also identifies which features are correlated with different labels, such as high DOC samples or low DOC samples. Feature importance determined by the neural network can be interpreted in the same way, where positive feature importance values imply a direct relationship with DOC, and negative values imply an inverse relationship. All 180 features shared by the neural network and the indicator species methods exhibit the same feature-label correlations. Fig 2C shows how prediction performance of the neural network and random forest models change as the number of features included in the model increases from a minimum of 10 features to a maximum of 86 features. The order in which features were included in each subsequent prediction corresponds to the rank determined by each feature selection method, such that the highest ranked features were included first. Both models reach close to peak prediction performance with only 86 features.

One might expect that the most informative features for DOC prediction would be those with highest or lowest abundances within communities. To examine this expectation, Fig 3A shows a histogram of bacterial abundance of the consensus set for selected features compared to the histogram of bacterial abundance for the entire dataset. This shows that feature selection techniques are not biased towards selection of taxa with low or high abundance, but rather the consensus set of taxa selected by random forest, neural network, and indicator species analysis had abundance levels mostly in the moderate range. Abundance values in the figure were determined for each taxon by taking the average number of reads over the entire set of samples. Fig 3B shows the distribution of prevalence of bacterial species of the consensus set of selected features, where prevalence was calculated as the frequency in which taxa were present in each sample. The distribution of prevalence of selected taxa shows that prevalence was not a crucial factor in selecting features for prediction of DOC.

**Fig 3.**
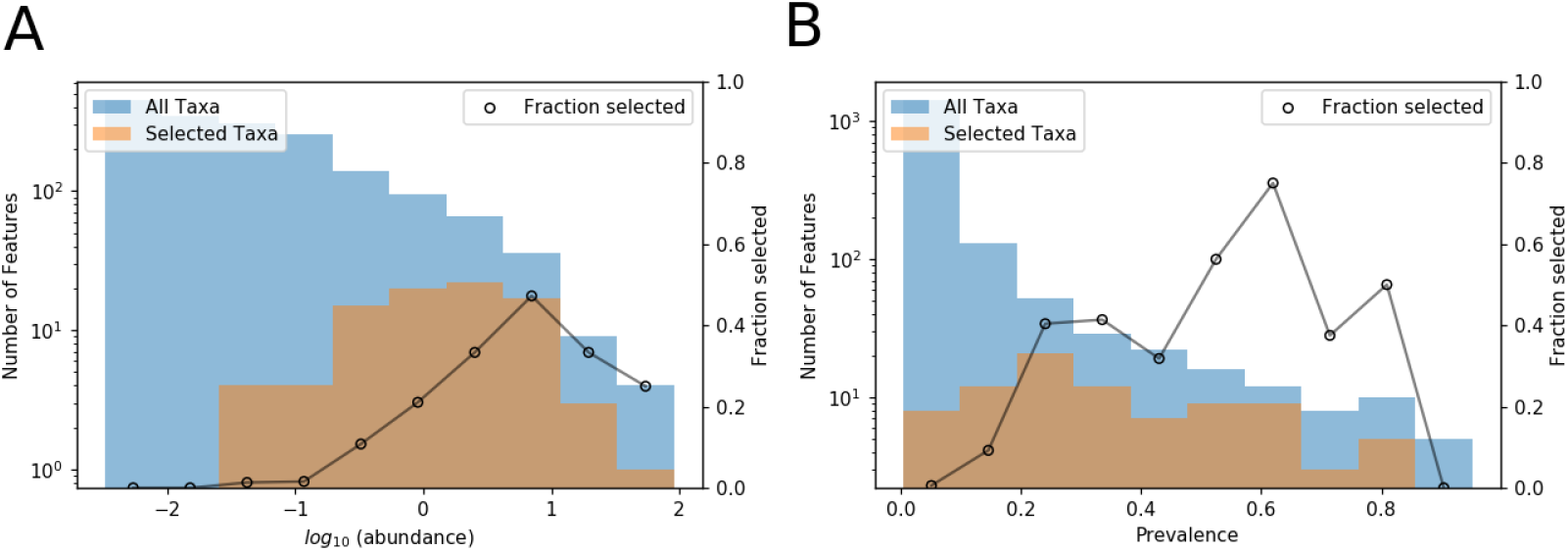
Distributions of bacterial abundance and prevalence of all taxa and the consensus set of taxa selected by all methods. (A) Histogram of abundance of taxa in the consensus set plotted over a histogram of abundance of all taxa in the data set. Abundance was calculated as the average number of taxa over the entire sample set. (B) Histogram of prevalence of taxa in the consensus set plotted over a histogram of prevalence of all taxa in the data set. Prevalence was calculated based on how frequently taxa were present in each sample.

To test generality of the above results, we determined the Pearson’s correlation coefficient for testing data under 50 randomly generated permutations of training and testing data with roughly 260 training samples and 50 test samples (exact sample sizes varied between 254 and 262 samples for training data and between 46 and 54 samples for test data due to variations in the number of replicates per experimental condition). Fig 4 shows histograms of test performance of the neural network model and the random forest model using the full feature set (Fig 4A,C) and the reduced feature set (Fig 4B,D). While the neural network model performed better using the reduced set of 86 features (two tailed t-test, *P* = .047), the distribution of prediction errors using the random forest model with the reduced feature set was not significantly different (two tailed t-test, *P* = .98). The neural network model produced greater prediction accuracy using the reduced feature set on 70% of test samples, and the random forest model yielded greater prediction accuracy on 48% of test samples. The random forest model significantly outperformed the neural network model with the full feature set (two tailed t-test, *P* < 0.001) but only marginally so with the reduced feature set (two tailed t-test, *P* = 0.11).

**Fig 4.**
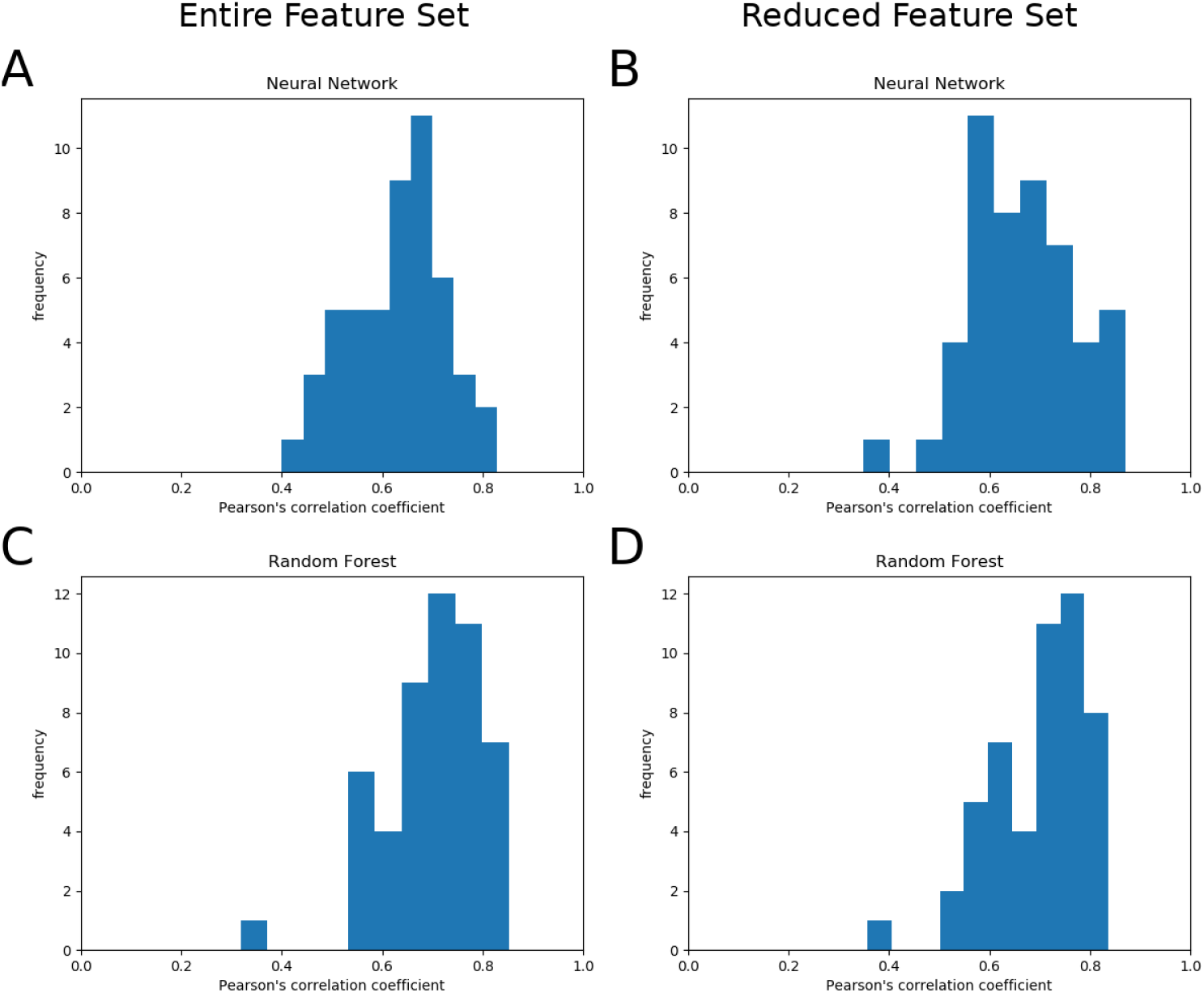
Distribution of prediction errors for 50 different permutations of training and testing data. (A) Distribution of Pearson’s correlation coefficients on test data performance using the neural network model without feature reduction. Mean R value = .627, standard deviation = .097. (B) Distribution of Pearson’s correlation coefficients on test data performance using the neural network model with the reduced feature set. Mean R value = .668, standard deviation = .103. (C) Distribution of Pearson’s correlation coefficients on test data performance using the random forest model without feature reduction. Mean R value = .699, standard deviation = .100. (D) Distribution of Pearson’s correlation coefficients on test data performance using the random forest model with the reduced feature set. Mean R value = .700, standard deviation = .095. For these permutations, feature reduction improved neural network prediction performance (two tailed t-test, *P* = 0.047), and random forest outperformed neural network with the full feature set (two tailed t-test, *P* < 0.001) and with the reduced feature set (two tailed t-test, *P* = 0.11).

To investigate how sample size affects model performance, prediction performance of the neural network and random forest regression models was measured with an increasing number of samples included in the training set (Fig 5). The random forest model consistently outperformed the neural network over the range of training data sample sizes, with more accurate predictions and less variability in prediction performance. Model performance of either method reaches near optimal levels after inclusion of only half of the training set or 150 training samples. Although variability in prediction performance continued to decrease as the fraction of training data increased, these results suggest that future experiments could be conducted with lower sample sizes without sacrificing model performance.

**Fig 5.**
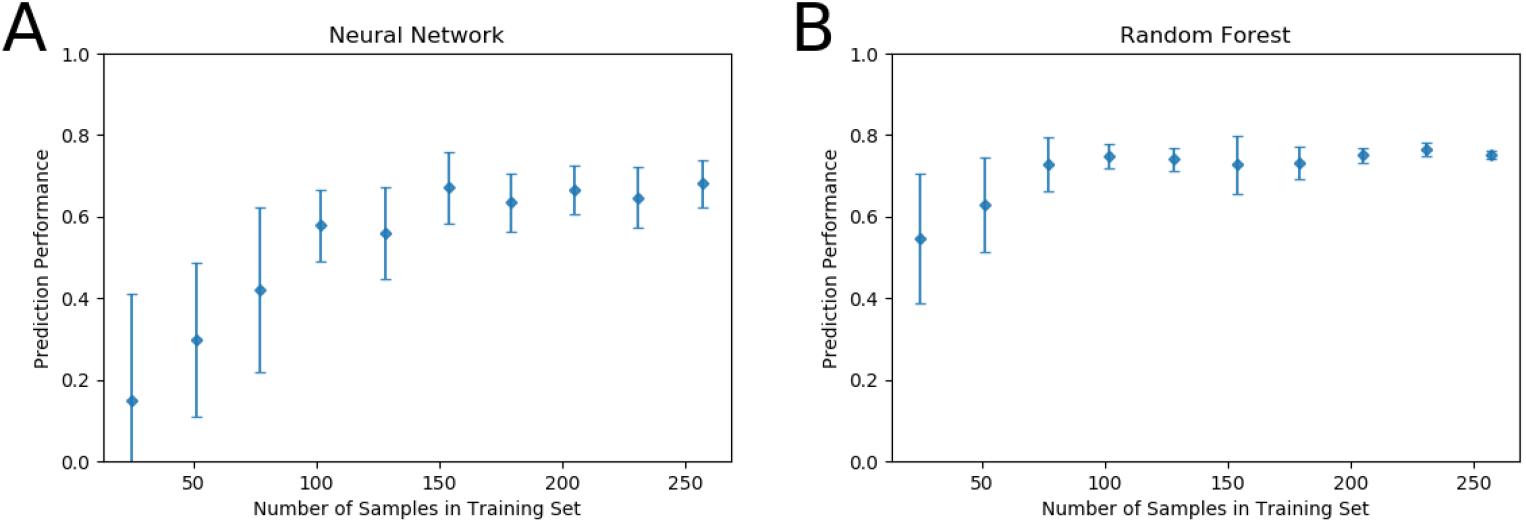
Sensitivity analysis of model prediction performance as the fraction of the total training data set (n=257) increases. Performance was measured using the average Pearson’s correlation coefficient after training over 10 random samplings of a fraction of the data set, with error bars representing 1 standard deviation from the mean. (A) Prediction performance on fixed testing data by the neural network model. (B) Prediction performance on fixed testing data by the random forest model.

## Discussion

While random forest outperformed the neural network for prediction tasks in this study, both methods can be used to predict DOC entirely from microbial community profiles and to provide measures of feature importance. The random forest method is relatively easy to implement, and performs well with little adjustment to model hyper-parameters. Sensitivity analyses with the data set in this study (Fig 5) shows that the random forest model is less sensitive to sample size of the training data set, which makes random forest an attractive machine learning model for analysis of microbiome data. A benefit of the neural network model is that it provides more easily interpreted results for feature selection, which include the direction in which taxa affect environmental variables. The site correlations determined by the neural network and indicator taxa analysis show perfect agreement in sign among the entire set of taxa. Furthermore, because ground truth for which taxa drive changes in environmental variables is not known, the joint set of selected features from random forest, neural network, and indicator taxa approaches provides greater confidence than the set from one method alone (feature selection results are included in the supporting information S5 Dataset).

Machine learning approaches for analyzing microbiome data have proven successful in applications such as forensics, medicine, and agroecology [26–28]. Recently, machine learning algorithms such as random forest and K-means clustering have successfully determined the postmortem interval (PMI) using postmortem skin microbiome [26]. In medicine, machine learning models such as random forest have been used for identification of gut microbiomes associated with irritable bowel syndrome in pediatric patients [27]. In another study focusing on soil microbiomes, a random forest model was applied to predict crop yields from soil microbiome composition [28]. With increasing access to machine learning software and high-dimensional microbiome data, machine learning is emerging as a powerful tool for understanding how microbial communities affect their environment.

Although there are several examples of platforms that facilitate use of machine learning techniques with microbial community data, our platform provides several unique options that make it more accessible and useful for microbial ecologists. QIIME [29] includes the “sample classifier” plugin [9], which provides access to a host of Sciklt-learn [21] implemented machine learning classification and regression models for use with microbiome data. Although the sample classifier QIIME plugin includes hyper-parameter optimization and feature selection of important bacterial taxa, it does not provide insight into directional relationships between bacterial taxa and target variables. Moreover, the sample classifier plugin is not set up to provide combined feature selection results determined from different machine learning methods, and feature selection is not determined using different permutations of the training data. MetAML [7] is another available software for implementing machine learning methods with microbiome data, but the methods are implemented exclusively for classification problems. For implementation of a neural network regression model with microbial abundance data, Neuroet [10] provides a simple GUI that can be used to train and test a single-layer, feed-forward neural network. Neuroet includes a procedure to optimize neural network architecture and identify important features for predicting model output, though optimization of hyper-parameters such as learning rate and the regularization coefficient is not available. While these platforms achieve a similar goal of applying machine learning techniques to microbiome data, no existing software packages include both neural network and random forest models and most do not provide insight into correlations between features and target variables. To provide the most confident set of important taxa, our tool produces the joint set of selected features from indicator species analysis, random forest, and neural network approaches. To aid in experimental design, our tool also provides a built-in tool for analyzing model sensitivity to experiment sample sizes.

Machine learning models offer the ability to determine hypothetical microbial communities that could promote increased levels of DOC. Enhancing carbon sequestration in soil is a strategy to combat climate change, as sequestration has the potential to offset fossil-fuel emissions by 0.4 to 1.3 gigatons (5 to 15 percent) of atmospheric carbon per year [14]. Under the assumption that a trained machine learning model has learned a general relationship between microbial abundance and DOC, we can use the model to determine a hypothetical microbial community that could potentially maximize DOC. In consideration of this task, the random forest and neural network models are markedly different. Although the random forest model has been at least as good as the neural network model to predict DOC levels that lie within the range of the previous training data, the random forest model is restricted by its formulation to a finite set of values corresponding to leaf nodes of decision trees. As a result, the random forest model is incapable of predicting values outside of the range presented in the training data. Conversely, the neural network model could in principle extrapolate to make predictions outside of the range present in the training data, which would enable specification of hypothetical microbial communities predicted to increase DOC beyond empirically observed levels. Furthermore, because the feature importance vector, **s**, produced by the neural network model is calculated as the gradient of the model output with respect to weights applied to features, **s** provides a potential direction in which features could be adjusted to increase levels of soluble carbon.

Fig 6A shows how the trained neural network model predicts responses to changes in microbial communities. In this simulation, communities (a) and (b) were initialized as the specific communities x_*a*_ and x_*b*_ that had the highest and lowest DOC and then adjusted in the direction defined by the feature importance vector according to x_new_ = x + *α*s, where *α* denotes the magnitude of the perturbation made to the community. The dashed trajectories represent DOC predictions made from simulated communities also initialized at the highest and lowest DOC, but with perturbations in random directions generated from a zero mean multivariate Gaussian distribution scaled by magnitude *α*. As the microbial community profiles were adjusted in the direction of the gradient determined by the neural network, the level of predicted DOC increased (see communities (a) and (b) in Fig 6A). When the same initial communities were adjusted randomly, predicted DOC never exceeded DOC predictions determined from communities x_*a*_ and x_*b*_ (see dashed blue and orange lines stemming from the same initial values as in communities x_*a*_ and x_*b*_). For the neural network model, community (a) results in predicted DOC levels that exceed the greatest DOC prediction from the training set, thus generating testable hypotheses to supplement communities to increase dissolved organic carbon. When the same simulated microbial communities were analyzed on a trained random forest model (Fig 6B), the model predicted a similar trend towards increasing DOC for community (b). Due to the nature of the algorithm, however, the level of DOC predicted by the random forest model could never exceed that of community (a). Simulation results using either model suggest that simulated communities informed by the trained neural network model are not random and produce theoretical microbiomes that could promote greater levels of carbon in soil, though future experiments are needed to test these designs and verify these predictions.

**Fig 6.**
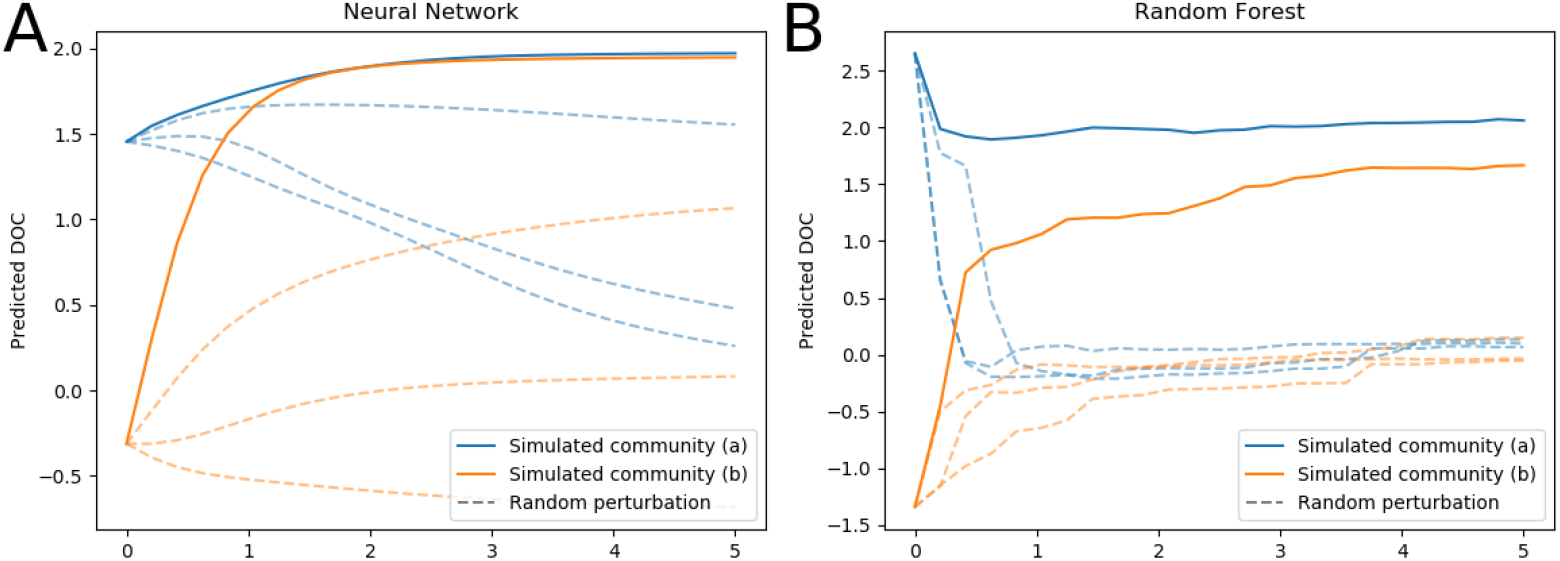
DOC predictions of trained machine learning models with synthesized microbial communities. Simulated communities (a) and (b) were specified by the training data communities with the highest and lowest DOC values, respectively. Each was then adjusted in the direction of the average gradient of maximum DOC increase determined by the neural network model, and each perturbation was scaled by magnitude *α*. Dashed lines stemming from the initial values of communities (a) and (b) represent DOC predictions from communities adjusted by a random vector with similar magnitude. (A) DOC prediction from hypothetical bacterial communities made by the neural network. (B) DOC prediction made by the random forest model with identical communities used in panel A.

Machine learning methods presented in this paper are intended to be easily applied to any data set that relates microbial communities to a scalar variable. To make this readily accessible, we have implemented all methods as a user-friendly platform available at https://github.com/MunskyGroup/thompson_et_al_plos_one_2019. For users without substantial knowledge of machine learning techniques, our tool enables application of machine learning regression models with optimized model parameters in a few lines of code. Tutorials for installing dependencies and using our machine learning tool can also be found on the GitHub repository. In this study, we applied machine learning approaches to elucidate the relationship between bacterial communities and carbon flow from plant litter decomposition by developing regression models to predict dissolved organic carbon (DOC) concentrations. For the dataset we analyzed from [17], a strong relationship exists between bacterial community composition and DOC abundance. Moreover, we found a consistent set of bacterial taxa identified by multiple methods – in this case neural network, random forest, and indicator species approaches.

With our platform, a table of feature selection results from random forest, neural network, and indicator species analysis is easily produced with a built-in feature selection function. Model sensitivity to sample sizes is also easily visualized using a built-in sensitivity analysis that plots prediction performance on testing data as the size of the training data set increases. The combination of machine learning tools and indicator species analysis reduced the feature set of 1709 taxa to 86 taxa, which is a critical step towards elucidating mechanistic relationships between microbial communities and environmental factors. Sensitivity analysis performed with the neural network and random forest models suggests that future studies could be performed with smaller sets of samples. Feature importance determined by the neural network could direct future studies by proposing microbial communities that enhance a functional outcome of interest, such as increased carbon flow into soil. In this context, the proposed machine learning tools provide a framework for designing experiments to further investigate how microbial communities function together to affect their environment.

## Acknowledgments

JT, RJ, JD and BM were supported by grant F255LANL2018 from the U.S. Department of Energy Office of Biological and Environmental Research, Genomic Science program. BM and JT were supported under award number R35GM124747 from the National Institute of General Medical Sciences of the National Institutes of Health. The funders had no role in study design, data collection and analysis, decision to publish, or preparation of the manuscript.

## Supporting information

**S1 Dataset OTU table.** The bacteria OTU table used for all results in the paper organized with samples as rows and features as columns.

**S2 Dataset Training data set.** Partition OTU table used for training machine learning models to produce Fig 1.

**S3 Dataset Testing data set.** Partition OTU table used for testing machine learning models to produce Fig 1.

**S4 Fig Monte Carlo simulation.** Monte Carlo simulation of the expected number of shared features after sampling from randomly organized sets of 1709 features.

**S5 Dataset Feature selection table.** A table with feature importance values for the consensus set of taxa sorted by the indicator species statistic.

