## Supplementary material for "Machine learning to predict microbial community functions: An analysis of dissolved organic carbon from litter decomposition": S4_Fig

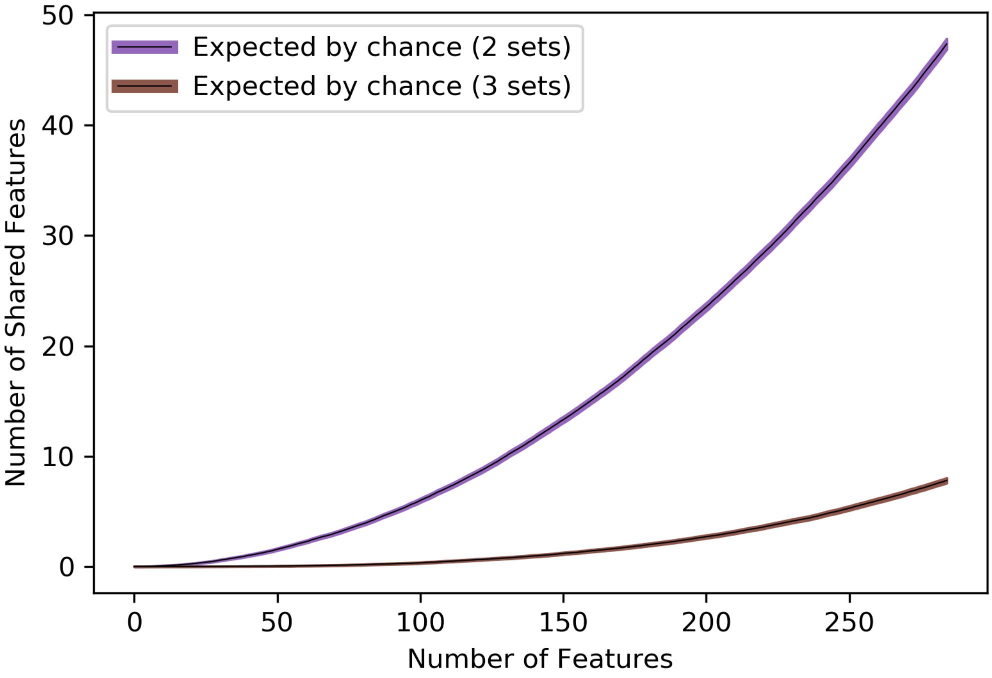


**S4 Fig. Monte Carlo simulation of the expected number of shared features after sampling from randomly organized sets of 1709 features.** Monte Carlo approach samples features from three randomly organized sets of 1709 features to count the number of features commonly selected in a pair of sets (purple) or within the intersection of all three sets (brown). Plotted curves show the mean and 99% confidence interval from 1,000 simulations as a function of the number of sampled features.
